# The role of cytonuclear interactions to plant adaptation across a *Populus* hybrid zone

**DOI:** 10.1101/2025.05.13.653687

**Authors:** Michelle Zavala-Paez, Brianna Sutara, Stephen Keller, Jason Holliday, Matthew C. Fitzpatrick, Jill Hamilton

**Affiliations:** Pennsylvania State University, University Park, PA, USA 16803; University of Vermont, Burlington, VT, USA, 05405; Virginia Tech, Blacksburg, VA, USA, 24061; University of Maryland Center for Environmental Science, Frostburg, MD, USA, 21532

**Keywords:** chloroplast, nuclear, co-introgression, hybridization, adaptation, physiology

## Abstract

Co-adaptation of cytoplasmic and nuclear genomes are critical to physiological function for many species. Despite this understanding, hybridization can disrupt co-adaptation leading to a mismatch between maternally-inherited cytoplasmic genomes and biparentally inherited nuclear genomes. Few studies have examined the consequences of cytonuclear interactions to physiological function across environments. Here, we quantify the degree of co-introgression between chloroplast and nuclear-chloroplast (N-cp) genes across repeated hybrid zones and its consequences to physiological function across environments. We use whole-genome resequencing and common garden experiments with clonally replicated genotypes sampled across the natural hybrid zone between *Populus trichocarpa* and *P. balsamifera*. We use geographic clines to test for co-introgression of the chloroplast genome with N-cp and non-interacting nuclear genes. Co-introgression of chloroplast and N-cp genes was limited although contact zone-specific patterns suggest that local environments may influence co-introgression. Combining ancestry estimates with phenotypic data across common gardens revealed that mismatches between chloroplast and nuclear ancestry can influence physiological performance, but the strength and direction of these effects vary depending on the environment. Overall, this study highlights the importance of cytonuclear interactions to adaptation, and the role of environment in modifying the effect of those interactions.

## Introduction

Organelle and nuclear genomes function together to maintain key developmental and physiological processes critical to adaptation [1–4]. In plants, these interactions are critical, as photosynthesis and respiration require coordination of nuclear-encoded proteins alongside the chloroplast and mitochondrial genomes [5–7]. Cytonuclear coadaptation predicts that nuclear genes involved in cytonuclear interactions will co-evolve with cytoplasmic genes to optimize plant function and fitness [8–10]. However, as cytoplasmic genomes evolve at a different rate relative to the nuclear genome, compensatory changes in nuclear-encoded genes that interact with cytoplasmic genomes are often required to maintain function [8–10]. This promotes positive intergenomic epistasis and is considered a major force shaping cytonuclear interactions [2,11]. Recombination between sister species via hybridization can also disrupt co-adaptation leading to a mismatch in ancestry with impacts to plant fitness [8,12–15]. Moreover, environmental gradients may favor specific combinations of cytoplasmic genes and their interacting nuclear genes that optimize physiological performance under local conditions [16–18]. Thus, understanding how mutation and recombination impact cytonuclear interactions across environments will be critical to ensuring optimized plant function in response to environmental change.

Despite the importance of co-adaptation between cytoplasmic and nuclear genomes to plant function, uniparental inheritance of the cytoplasmic genome with biparental inheritance of the nuclear genome has frequently led to a mismatch in genomic ancestry where sister species hybridize [2,15,19]. As a result, hybrids may have a cytoplasmic genome from one species while exhibiting a mixed nuclear background resulting in negative intergenomic epistasis for co-adapted genes [2,15,19,20]. Indeed, where previously isolated lineages have come into secondary contact accumulated differences have contributed to genetic incompatibilities with negative impacts to fitness [21,22]. Experimental studies have shown that such mismatches can impair metabolic processes, reduce photosynthetic capacity, contribute to chlorosis, and ultimately decrease fitness [8,12–15]. Given these potential impacts, understanding how and when hybridization influences cytonuclear interactions and their genetic and phenotypic consequences across environments will be critical for predicting plant performance.

Despite the potential fitness consequences of cytonuclear mismatch for plants, cytoplasmic genomes frequently introgress across species boundaries and this introgression may play a role in adaptation to new environments [23–25]. Cytoplasmic introgression has been shown to enhance organellar function [15,26] and can provide a mechanism to mitigate the potential fitness consequences of high cytoplasmic mutational load [23,27,28]. However, because nuclear and cytoplasmic genomes interact, the benefits may depend on co-introgression of cytoplasmic genomes alongside their nuclear-encoded interacting genes [23,29]. Theory predicts that such genes that interact across the cytoplasm and nuclear genome should be more likely to introgress alongside cytoplasmic genomes, irrespective of nuclear genes without such interactions (hereafter, non-interacting nuclear genes). [13,23,30]. Since cytonuclear interactions can be shaped by local environmental conditions [16,17] evaluating patterns of co-introgression across multiple contact zones in a hybrid zone offers a unique opportunity to assess how environment influences these interactions. Despite this understanding, it remains unclear whether cytoplasmic genomes and their interacting nuclear genes co-introgress more frequently than non-interacting nuclear genes across repeated zones of natural hybridization.

*Populus* is an ideal system for studying cytonuclear interactions as repeated hybridization between sister species has led to frequent evidence of cytoplasmic introgression, resulting in natural variation in cytonuclear combinations [25,31–33]. Here, we focus on the widespread natural hybrid zone between *P. trichocarpa* and *P. balsamifera*, which spans a steep maritime-continental climatic gradient characterized by extensive nuclear gene flow and the formation of advanced-generation hybrids [34–36]. Within this hybrid zone, species-specific chloroplast haplotypes are divergent, but substantial interspecific gene flow has resulted in a spectrum of individuals with varying degrees of nuclear and chloroplast ancestries [31,34,37]. This variation provides a unique opportunity to evaluate co-adaptation between the nuclear and chloroplast genome, specifically by testing whether nuclear-encoded genes that interact with the chloroplast genome (hereafter, N-cp genes) and chloroplast genome co-introgress and by quantifying how varying cytonuclear combinations affect traits important to plant physiological function. Despite recognition of the evolutionary significance of hybridization in *Populus* and the extensive use of *Populus* hybrids in restoration and bioenergy, the role of cytonuclear interactions to adaptation remains largely untested.

Using whole-genome resequencing data from *Populus* genotypes sampled across repeated natural hybrid zones between *P. trichocarpa* x *P. balsamifera*, we test for patterns of co-introgression between N-cp genes and the chloroplast genome and evaluate the phenotypic effects of cytonuclear interactions to physiological trait variation across environments. Given its compact size, conserved gene content, and central role in photosynthesis, we use the chloroplast genome as our focal cytoplasmic genome to evaluate patterns of co-introgression, focusing specifically on nuclear–chloroplast interactions [25]. We address three main questions: (1) Is there evidence of chloroplast introgression across multiple contact zones within the hybrid zone between *P. trichocarpa* and *P. balsamifera*, and what influence does environment have on the extent or direction of chloroplast introgression? Given previous reports of extensive gene flow between *Populus* species, we predict that introgression of the chloroplast genome may be environmentally dependent (2) Do geographic cline comparisons between N-cp genes and the chloroplast alongside non-interacting nuclear genes indicate co-introgression for genes critical to adaptation? If cytonuclear coadaptation is maintained by selection, then N-cp genes should exhibit clinal variation that parallels chloroplast ancestry indicative of co-introgression. In contrast, non-interacting nuclear genes are expected to vary independently, exhibiting clinal patterns independent of the chloroplast genome (3) Do varying cytonuclear combinations influence physiological traits critical for adaptation, and if so, do these effects vary across environments? We expect that cytonuclear discordance will reduce physiological performance relative to matched combinations due to disruption of co-adapted gene complexes. By addressing these questions, our study provides new insights into how cytonuclear interactions shape adaptation, emphasizing the role of environmental selection in structuring patterns of co-introgression and trait variation in *Populus*.

## Methods

### Plant material sampling, genomic library preparation and variant calling

In January 2020, dormant vegetative cuttings were sampled from 574 individual *Populus* genotypes across seven latitudinally distributed contact zones spanning the natural hybrid zone between *Populus trichocarpa* and *P. balsamifera* (Figure S1, Supplementary Table S1 and S2; full details described in Bolte et al. 2024). Cuttings were transported to Blacksburg, VA, USA where they were divided and propagated, creating a set of clonally replicated rooted *Populus* cuttings. Clones were maintained in the greenhouse at 24 °C daytime and 15.5 °C nighttime temperatures without the use of supplemental lighting (Full details described in Bolte et al. 2024).

Following vegetative flush, 100 mg of fresh leaf tissue was collected from rooted cuttings for DNA extraction and whole genome resequencing. Methods regarding DNA extraction, genomic library preparation, and subsequent quality control and filtering associated with whole-genome resequencing of the nuclear genome are available in Bolte et al. (2024). Briefly, genomic libraries were sequenced on a S4 flow cell in 2x150bp format on an Illumina NovaSeq 6000 instrument with 64 samples per lane. For each genotype, Illumina reads were mapped to the *P. trichocarpa* reference genome (v4.0) and stored as SAM files. SAM files were converted to BAM format with SAMtools [38]. Individual gVCF files were created using the Genome Analysis Toolkit (v3.7) Haplotype Caller algorithm and merged into a single VCF file using the GATKGenotype GVCFs function, with the raw VCF file containing 94,540,211 variant calls. The raw VCF file was filtered for map quality (MQ < 40.00), elevated strand bias (FS > 40.000, SOR > 3.0), differential map quality between reference and alternative alleles (MQRankSum < - 12.500), positional bias between reference and alternate alleles (ReadPosRankSum < -8.000), and depth of coverage (QD < 2.0), which results in a dataset comprised of 45,690,186 variant calls. INDELS were removed along with SNPs with ≥ 2 alternate alleles, and variants were further filtered based on a minimum allele frequency (MAF < 0.05) and no missing data resulting in 4,497,721 biallelic SNPs for subsequent analysis.

For the chloroplast genome, variant calling was performed using the paired-end Illumina reads for all 574 samples within bcftools 1.14 [39]. The *P. trichocarpa* chloroplast genome (NC_009143.1) was used as reference; however, before mapping one inverted repeat region was manually removed to avoid misalignment. Reads were aligned to the modified *P. trichocarpa* chloroplast genome using Bowtie and stored as SAM files using Samtools 1.15 [38]. Subsequent variant calling was performed on the individual sorted BAM files assuming a haploid model (no heterozygosity) without Hardy-Weinberg Equilibrium expectations. The VCF file was filtered with a minimum variant quality threshold (QUAL) of 30, maximum missing data of 0.05 with indels were removed. A total of 1,879 variants were kept after filtering for subsequent analysis.

### Chloroplast genome assembly, chlorotype identification, and divergence

Chloroplast genomes were assembled using NOVOPlasty [40], with assembly parameters detailed in Supplementary Methods. Genomes were annotated in Geneious v7.1.4 [41] using *P. trichocarpa* (NC_009143.1) as a reference, with start and stop codons manually verified. Of the 574 samples, 573 chloroplast genomes were successfully assembled, with one sample (GPR-14_S50_L001) failing due to low coverage (<10×), and 10 individuals removed due to assembly errors in the small single-copy region (SSC).

To infer species-specific chloroplast ancestry, we used 563 chloroplast genomes from our collection alongside 58 publicly available *Populus* spp. chloroplast genomes and *Salix interior* and *S. tetrasperma* chloroplast genomes from GenBank (Supplementary Table S3). Chloroplast genomes were aligned using MAFFT v7.453 [42], and a maximum likelihood (ML) tree was constructed in IQ-tree 1.6.11. The best-fit evolutionary model (TVM+F+R2) was selected using ModelFinder [43] based on the lowest BIC score, and branch support was assessed with 1,000 bootstrap replicates. Ancestry was assigned based on species-specific clades using FigTree v1.4.13 (Supplementary Figure S2 and S3). To further validate phylogenetic inference within and among clades, BLAST searches of *matK* and *rbcL* genes were performed, confirming chloroplast ancestry when genotypes and reference sequences exhibited 99% sequence similarity.

To assess differences associated with the chloroplast genome between *P. trichocarpa* and *P. balsamifera*, species-specific SNPs were identified. Fixed differences were quantified using a custom script in R. Following this, SnpEff [44] was used to determine the frequency of non-synonymous and synonymous substitutions relative to the *P. trichocarpa* reference chloroplast genome to identify species-specific differences.

### Testing for the role of environment on chloroplast ancestry

Logistic regression was used to assess the relationship between environment and chloroplast ancestry across the six repeated contact zones between *P. trichocarpa* x *P. balsamifera*. We used climate averages associated with genotype origin (1961-1990) for six climatic variables previously associated with nuclear genetic structure [34]. These variables include temperature differential, a measure of continentality (temperature difference between mean warmest month temperature and mean coldest month temperature, TD), mean annual temperature (MAT), mean annual precipitation (MAP), climate moisture deficit (CMD), relative humidity (RH), and precipitation as snow (PAS) downloaded from climateNA V742 [45]. Firth’s Bias-Reduced Logistic Regression was used to account for small sample sizes and to address complete separation, which occurs when predictors perfectly separate outcomes [46,47]. Logistic regression models were fitted using the logistf function in R 4.3.1 to quantify the strength and direction of associations between chloroplast ancestry and environmental gradients [48].

### Geographic clines to compare patterns of introgression between chloroplast and nuclear-interacting genes

Geographic clines were compared between the chloroplast genome, nuclear-chloroplast (N-cp) genes, and non-interacting nuclear genes to test expectations for co-introgression across latitudinally-distributed contact zones. We selected 282 N-cp genes previously identified in *Arabidopsis thaliana* from the Cytonuclear Molecular Interactions Reference (CyMIRA) database [49] and identified their *Populus* orthologs using BLASTP (e-value < 10⁻⁵, minimum identity = 60%). Non-interacting nuclear genes were selected from 2,668 single-copy nuclear genes in *P. trichocarpa* after excluding all nuclear genes with known cytonuclear interactions listed in CyMIRA, resulting in 2,611 nuclear genes for use in downstream analysis.

Locus-specific ancestry for each of the N-cp and non-interacting nuclear genes was inferred using Loter [50], which assigns species-specific ancestry using phased reference panels of unadmixed parental species, estimating ancestry for linkage blocks within admixed individuals. Parental-type genotypes were identified previously from [51] based on admixture with Q > 0.98 and Q < 0.02 (for K=2) assigned as *P. trichcocarpa* (*n* =131) and *P. balsamifera* (*n* = 90), respectively. Local ancestry was assessed for 323 admixed individuals, and gene-specific ancestry values were extracted from the ancestry matrix using gene positions from the *P. trichocarpa* v4.1 reference genome, including a ±100 bp flanking region. For each admixed individual, locus-specific ancestry for each gene was calculated as the proportion of *P. trichocarpa* ancestry while chloroplast ancestry was assigned based on phylogenetic inference.

Geographic clines were modeled for chloroplast ancestry, 282 N-cp, and 2,611 non-interacting nuclear genes using the R package HZAR [52]. Cline parameters, including center and width (calculated as 1/maximum slope), were used to compare introgression across groups. For each of the latitudinally-distributed contact zones, geographic distance from the coast for each genotype was calculated using the Haversine formula and normalized between 0 and 1 for comparison (Supplementary Methods). Clines were modeled and compared for different scale and tail parameters with Akaike’s Information Criterion (AIC) used to select the best fit model (For details, see Supplementary Methods). Cline parameters were compared between N-cp genes, chloroplast genome and non-interacting nuclear genes to test expectations of co-introgression. Co-introgression was observed when N-cp gene cline parameters overlapped with the 95% confidence interval (CI) of the chloroplast cline parameters, but not with the 95% CI for non-interacting nuclear genes. To determine whether the pattern of introgression for N-cp genes differs significantly from either chloroplast or non-interacting nuclear genes, we fit two constrained models for each N-cp gene in which the cline center was fixed within the 95% CI of either the chloroplast or non-interacting nuclear genes cline center using HZAR [52]. We then performed likelihood ratio tests (LRTs; 1 df) to assess whether the best-fit model for each N-cp gene differed significantly from either constrained model.

### Common gardens experiments

A total of 574 unique *Populus* genotypes were clonally propagated and planted in March 2020 in two common gardens located in Blacksburg, VA (36°37′N, 80°09′W; 359 m) and Burlington, VT (44°26′N, 73°11′W; 85 m; Figure 1). Each garden was established using a randomized complete block design with three replicates per genotype, totaling 1,722 trees per garden. During summer 2022, physiological traits related to chloroplast function, stomatal conductance, water-use and nitrogen-use efficiency were evaluated across sites. To limit variation due to developmental stage [53], the first fully expanded leaf of the dominant shoot was sampled to assess stomatal conductance (g_sw_, mmol m⁻² s⁻¹) and chloroplast fluorescence parameters including quantum efficiency of photosystem II (ΦPSII), minimum fluorescence (Fs), maximum fluorescence (Fm), and electron transport rate (ETR, µmol m⁻² s⁻¹) using an LI-600 (LI-COR). Chloroplast fluorescence values represent the average of three measurements taken from non-overlapping regions per individual leaf. The second fully expanded leaf was sampled for carbon (δ¹³C, ‰) isotopes and nitrogen (N, %) content analyses. Leaves were dried at 60°C, homogenized using a TissueLyser II (Qiagen, Germany), and analyzed at the Central Appalachians Stable Isotope Facility (CASIF, University of Maryland Center for Environmental Science, USA).

**Figure 1.**
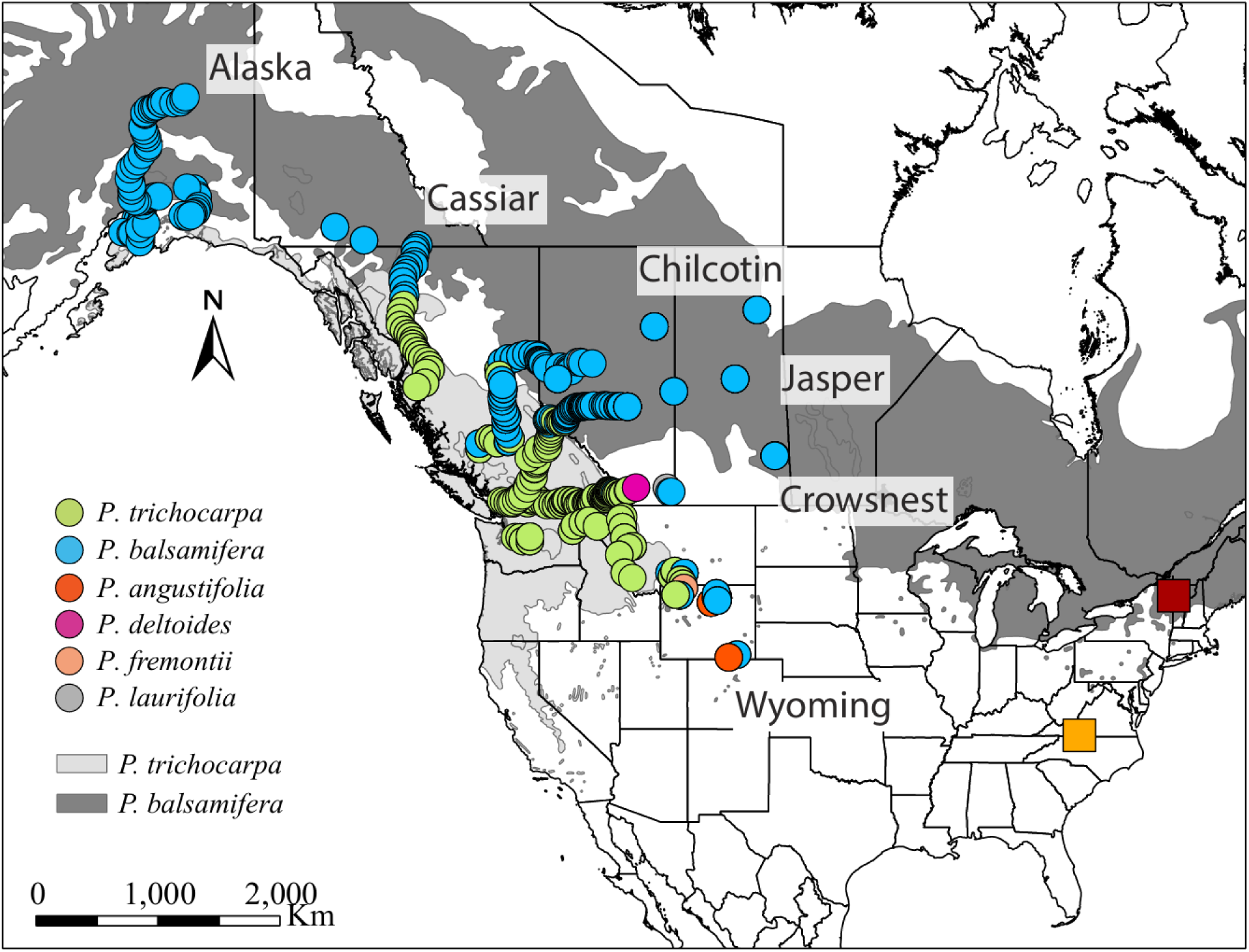
Genotypes were collected across six east-west contact zones within the natural hybrid zone between *P. trichocarpa* and *P. balsamifera*. Light and dark gray represent the distribution range of *P. trichocarpa* and P*. balsamifera*, respectively. Georeferenced sampling locations are color coded by chloroplast ancestry, *P. trichocarpa* (green), *P. balsamifera* (blue), *P. angustifolia* (orange), *P. fremontii* (peach), *P. deltoides* (purple), and *P. laurifolia* (gray). B) Clonally replicated genotypes were planted at the common gardens in Virginia Tech (VT, yellow square), and University of Vermont (UVM, red square).

### Heritability

To quantify the proportion of phenotypic variance explained by genetic differences among genotypes, we estimated broad-sense heritability (*H²)* for each trait across each common garden. Variance components, genetic variance (*V*_*G*_) and total phenotypic variance (*V*_*P*_), were estimated by fitting a linear mixed-effects model for each garden using equation 1:

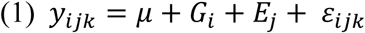

Where *y*_*ijk*_ is the observed trait value for the individual *k* in the genotype *i* and block *j*. *G*_*i*_ is the random effect of genotype representing the genetic effects (*V*_*G*_), *E*_*j*_ is the random effect of block representing environmental effects (*V*_*E*_) and *ɛ*_*ijk*_ is the residual error (*V*_*ɛ*_). Variance components from the linear mixed-effects model were extracted using VarCorr in R 4.3.1. *H²* was then calculated using equation 2:

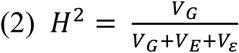

### Statistical analysis

To quantify the effect of the interaction between nuclear and chloroplast ancestry to physiological trait variation across environments, each trait was modeled using equation (3):

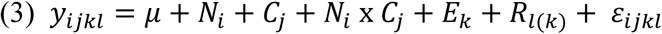

Where, *N*_*i*_ is the fixed effect of nuclear genome ancestry *i*, *C*_*j*_ is the fixed effect of chloroplast ancestry *j*, and *N*_*i*_ x *C*_*j*_ corresponds to the interaction between them. *E*_*k*_ is the fixed effect of common garden environment *k* accounting for environmental differences across common garden experiments. The random effect of block nested within garden *R*_*l*(*k*)_ was included to account for site-specific environmental variation within garden and *ɛ*_*ijkl*_ is the error term. A second model was tested to include the interaction between chloroplast ancestry and environment, as well as the three-way interaction among nuclear ancestry, chloroplast ancestry, and environment. However, these interactions were not significant, so the simplified model was used (Supplementary Methods). Models were fitted using the package lme4 [54] in R 4.3.1. For each model, normality of residual errors was visually assessed.

## Results

### Chloroplast ancestry and geographic distribution across the hybrid zone

Phylogenetic analysis of 563 genotypes indicated that 247 genotypes had a chlorotype associated with *P. trichocarpa* and 298 genotypes had the *P. balsamifera* chlorotype (Supplementary Figure S2). Of the remaining 18 genotypes, a BLAST comparison of *Populus* chloroplast genomes revealed varying ancestry with 99% similarity with either *P. laurifolia* (5), *P. deltoides* (1)*, P. fremontii* (4) or *P. angustifolia* (8) (Supplementary Figure S3). In addition, across repeated contact zones between *P. trichocarpa* and *P. balsamifera*, the distribution of chlorotypes varied. In the northern Alaska contact zone, *P. balsamifera* was the sole chlorotype observed indicating a likely chloroplast capture (Figure 1). Both chlorotypes were observed across all other contact zones; however, the *P. balsamifera* chlorotype extended across the entire geographic extent of the Chilcotin contact zone.

### Chloroplast genome structure and genetic variation

Chloroplast structure was highly conserved across individuals for both chlorotypes, with typical quadripartite organization (LSC, SSC, IRA, and IRB; Supplementary Figure S4). The chloroplast genome for both species contained 78 coding genes. A comparison between the chlorotypes of *P. trichocarpa* and *P. balsamifera* identified 53 species-specific differences (Supplementary Table S4). Of these, 16 were found within coding regions (Supplementary Figure S4), resulting in eight amino acid changes within the following chloroplast genes: *matK* (1), *ndh*D (1), *ndh*K (1), *rpo*A (1), *rpo*C2 (1), *rps*8 (1), and *ycf*1 (2). These results suggest that while gene content is conserved between chlorotypes, there are species-specific differences that may underlay functional variation.

We compared chloroplast and nuclear ancestry across contact zones. Notably, while individuals with the *P. balsamifera* chlorotype were observed across the full extent of nuclear genome ancestry, the *P. trichocarpa* chlorotype was restricted to individuals with 42% to 100% *P. trichocarpa* nuclear ancestry across all six contact zones (Figure 2A). This suggests that barriers to *P. balsamifera* chloroplast introgression may exist where the nuclear genome for *P. trichocarpa* is most prevalent.

**Figure 2.**
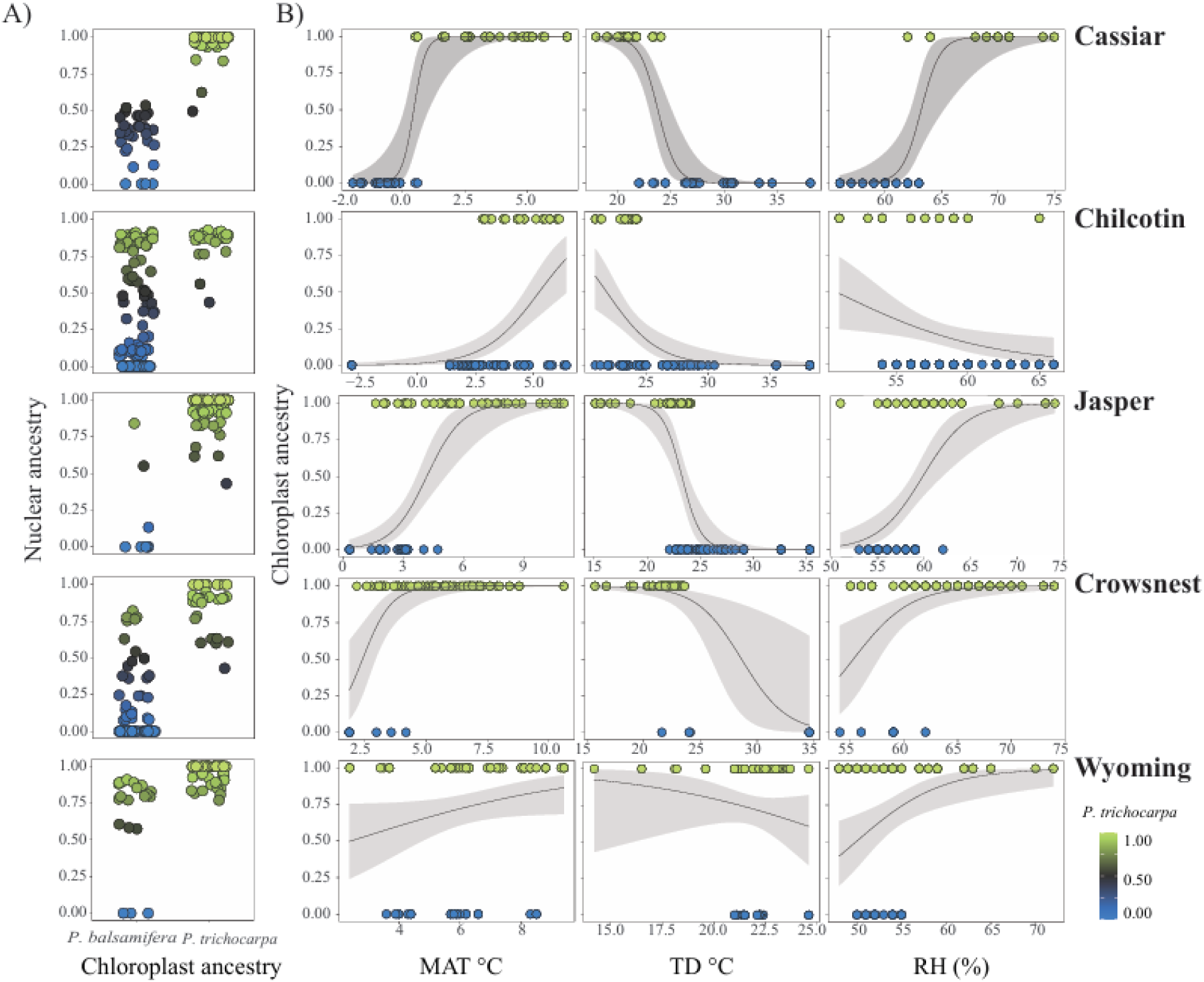
A) Chlorotype distribution (x-axis: *P. balsamifera* or *P. trichocarpa*) associated with the mean nuclear genome-wide ancestry (y-axis) per contact zone. Each point represents a genotype. Colors reflect nuclear genome-wide mean local ancestry of each genotype, ranging from *P. balsamifera* (blue) to *P. trichocarpa* (green), with admixed ancestry in grey. B) Relationship between chloroplast ancestry with the mean annual temperature (MAT), continentality (TD), and relative humidity (RH). Chloroplast ancestry (0 = *P. balsamifera*, 1 = *P. trichocarpa*) is represented as a function of a climate variable. Data points are color-coded by chloroplast ancestry: *P. balsamifera* in blue and *P. trichocarpa* in green. The gray line represents the predicted probabilities of chloroplast ancestry based on the fitted logistic regression, with the shaded blue area indicating 95% confidence intervals for the predictions.

### Differential influence of climate on chloroplast ancestry across repeated contact zones

Logistic regression models revealed that chloroplast ancestry was strongly associated with climatic gradients, although the strength and direction of these associations varied across contact zones (Figure 2B). Overall, mean annual temperature (MAT), continentality (TD), and relative humidity (RH) were the strongest predictors for chlorotype across the six contact zones with *P. trichocarpa* chlorotype more common in warmer, wetter environments and less continental environments relative to the *P*. *balsamifera* chlorotype (Supplementary Table S5). In the Cassiar contact zone, sharp transitions associated with MAT (*slope* = 4.23 per °C, *p* < 0.001) and RH (*slope* = 1.18 per %, *p* < 0.001) were observed, with *P. balsamifera* chlorotypes restricted to colder (–0.2 to 0.6 °C MAT) and drier (56–63% RH) environments, while *P. trichocarpa* chlorotypes occupied warmer (0.6–6.6 °C MAT) and increasingly humid (62–75% RH) regions (Figure 2B). The probability of *P. trichocarpa* presence decreased with TD (*slope* = –1.17 per °C, *p* < 0.001) and CMD (*slope* = –0.02 per mm, *p* < 0.001) but increased with MAP (*slope* = 0.02 per mm, *p* < 0.001) and PAS (*slope* = 0.01 per mm, *p* < 0.001) suggesting that climate is likely a selective force shaping the chloroplast distribution across the Cassiar contact zone.

Similar associations were observed in the Chilcotin, Jasper and Crowsnest contact zones, where *P. trichocarpa* chloroplast presence increased with warmer (*slope* = 0.85–1.69 per °C, *p* < 0.001) and less continental (*slope* = –0.48 to –1.12 per °C, *p* < 0.001) environments, although the strength of these associations was lower with respect of Cassiar (Supplementary Table S5). Additionally, RH was positively associated with *P. trichocarpa* presence in Crowsnest and Jasper (*slope* = 0.37–0.41 per %, *p* < 0.001), but had a negative effect in Chilcotin (–0.19 per %, *p* < 0.01).

In the Wyoming contact zone, temperature and continentality did not influence chloroplast ancestry distribution, but the *P. trichocarpa* chloroplast increased with RH (*slope =* 0.21 per %, *p* < 0.001), MAP (*slope* -0.01 per mm, *p* < 0.001), and PAS (*slope* = 0.01 per mm, *p* < 0.01), pointing towards the influence of moisture rather than temperature as selective force within this contact zone. Overall, these results indicate that chloroplast ancestry is influenced by climate, but the strength and direction of these associations vary by contact zone.

### Directional introgression of the *P. balsamifera* chlorotype with limited evidence of co-introgression between chloroplast and nuclear-interacting genes

A comparison of geographic clines revealed directional chloroplast introgression, with *P. balsamifera* chloroplasts moving into the *P. trichocarpa* geographic range (Figure 3). In Alaska, there was a chloroplast capture for the *P. balsamifera* chlorotype, therefore no clinal gradients were assessed. In the Cassiar contact zone, a steep, localized transition between chlorotypes was observed (*w* = 0.05, *c* = 0.43, Figure 3A, Supplementary Table S6) suggesting potential barriers to chloroplast introgression. Interestingly, 12 of 234 N-cp genes co-introgressed with the chloroplast within this contact zone (Figure 3B). These included genes involved in RNA processing, proteolysis, photosystem I, ribosomal proteins, and transfer RNAs (Supplementary Table S7) suggesting that where co-introgression was observed it included those N-cp genes involved in essential chloroplast function.

**Figure 3.**
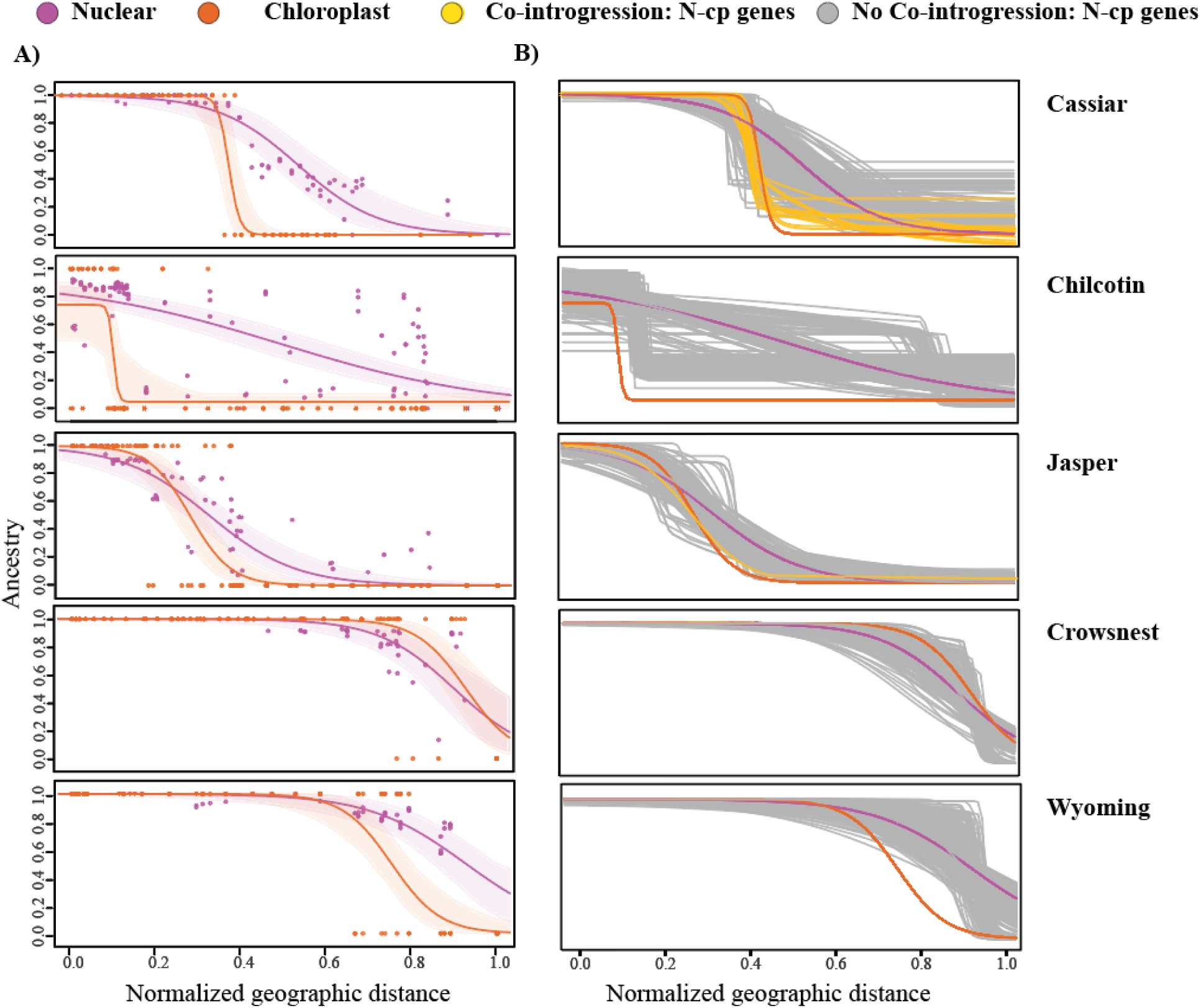
A) Maximum likelihood geographic clines for chloroplast ancestry (orange line) and mean non-interacting nuclear genes (purple line). The x-axis represents the scaled distance from the coast for each of the five contact zone. The y-axis represents ancestry proportion of ancestry associated with either *P. trichocarpa* (1) or *P. balsamifera* (0). Lines of best fit model were estimated using HZAR (Derryberry *et al.*, 2014) and the 95% confidence interval are shown. Dots represent an individual genotype mean ancestry for non-interacting nuclear genes (purple) and chloroplast ancestry (orange). B) Co-introgression tests for nuclear-chloroplast (N-cp) genes and chloroplast across five latitudinal contact zones. Geographic clines for N-cp genes (yellow and gray lines), chloroplast ancestry (orange lines), and mean non-interacting nuclear genes (purple lines) were estimated using HZAR. N-cp genes that co-introgress with the chloroplast genome are shown in yellow, while those that do not appear to co-introgress with the chloroplast genome are denoted in grey.

Across the Chilcotin and Jasper contact zones, chloroplast clines were wider (*w* = 0.04, 0.19, Figure 3A, Supplementary Table S6) consistent with weak barriers to gene flow. Cline centers were geographically located within the range of distribution of *P. trichocarpa* (*c* = 0.10, 0.27, Supplementary Table S6), suggesting introgression of the *P. balsamifera* chlorotype. In general, N-cp genes did not exhibit concordant cline center or widths with the chloroplast genome, except for PNSB3 in Jasper, suggesting a lack of co-introgression between chloroplast and N-cp genes (Figure 3B). Both the Crowsnest and Wyoming contact zones exhibited the widest chloroplast clines (*w* = 0.24, 0.23; *c* = 0.93, 0.75), with the transition between chlorotypes occurring towards the geographic interior of the hybrid zone (*c* = 0.93, 0.75). There was no evidence of co-introgression between the chloroplast and N-cp genes, suggesting that introgression of N-cp genes is independent of the chloroplast within these contact zones.

### Cytonuclear interactions influence physiological traits

Traits related to chloroplast function (ΦPSII, Fm, Fs, and ETR) exhibited low heritability values (0.00 – 0.08, Table 1) regardless of the planting environment. For the Vermont and Virginia common garden, water-use efficiency (δ^13^C, 0.21 - 028), nitrogen content (0.13 – 0.22) and g_sw_ (0.17 – 0.32) had higher heritability estimates, indicating increased genetic contribution to variation in these traits independent of the planting environment.

**Table 1.** Results of linear mixed-effects models testing the effects of nuclear ancestry (N), chloroplast ancestry (C), common garden environment (E), and cytonuclear interactions (N x C) on physiological traits in hybrid *Populus* genotypes. Nuclear ancestry represents the proportion of *P. trichocarpa* nuclear background. Chloroplast ancestry is represented as a factor, with *P. trichocarpa* chlorotype as reference. Garden environment is a factor, with UVM (University of Vermont) as reference. Slopes of the fixed effects (G, C, E, G×C) indicate the magnitude and direction of the effect on trait values; bold values indicate statistically significant effects (*p* < 0.05). Marginal R^2^ indicates variance explained by fixed effects alone, and conditional R^2^ includes both fixed and random effects.

| Trait | Heritability (H<sup>2</sup>) | Nuclear ancestry (N) | Chloroplast ancestry (C) | Common garden environment (E) | N x C | Marginal R<sup>2</sup> | Conditional R<sup>2</sup> |
| --- | --- | --- | --- | --- | --- | --- | --- |
| Quantum efficiency in light in PSII (ΦPSII) | 0.01 - 0.04 | 0.06 | **0.09** | -0.04 | **-0.11** | 0.03 | 0.20 |
| Maximum fluorescence in light (Fm) | 0.00 - 0.08 | 23.10 | **48.21** | -4.52 | **-54.31** | 0.01 | 0.16 |
| Minimum fluorescence in light (Fs) | 0.00 - 0.08 | -4.99 | 0.03 | **8.03** | 1.33 | 0.03 | 0.06 |
| Electron transport rate (ETR, μmol m<sup>-2</sup> s<sup>-1</sup>) | 0.00 - 0.01 | -19.88 | -7.68 | 28.63 | 8.92 | NS | NS |
| Carbon isotopes (δ<sup>13</sup>C, ‰) | 0.27 - 0.28 | 0.49 | -0.12 | -0.04 | -0.09 | NS | NS |
| Nitrogen content (N, %) | 0.13 - 0.22 | -0.28 | -0.26 | **-0.24** | **0.43** | 0.03 | 0.05 |
| Stomatal conductance (g<sub>sw</sub>, m<sup>-2</sup> s<sup>-1</sup>) | 0.17 - 0.32 | **-0.17** | 0.02 | **-0.26** | 0.02 | 0.42 | 0.47 |

Cytonuclear interactions significantly affected efficiency of photosystem II (ΦPSII) and maximum fluorescence in light (Fm), with mismatched nuclear and chloroplast ancestries reducing the efficiency of light absorbance (*p* < 0.01, *slope*=-0.11 and -52.65, respectively) (Figure 4A). Individuals with a *P*. *balsamifera* chlorotype had a slight efficiency advantage where nuclear ancestry was intermediate (50% P. trichocarpa ancestry). However, as *P. trichocarpa* nuclear ancestry increased beyond ∼75%, individuals genotypes with a mismatched chloroplast ancestry (*P. balsamifera* chloroplast) exhibited reduced efficiency. Interestingly, the negative effect of the interaction between nuclear and chloroplast ancestry on ΦPSII and Fm was similar across the two common gardens (*p >* 0.05, Figure 4A). We also measured Fs, ETR, and δ¹³C and noted no effect of either nuclear genome ancestry, chloroplast genome ancestry, or their interactions on phenotypic variation (Table1, Supplementary Figure S6). Overall, these results suggest that mismatched nuclear-chloroplast combinations can reduce performance for key photosynthetic traits across environments.

**Figure 4.**
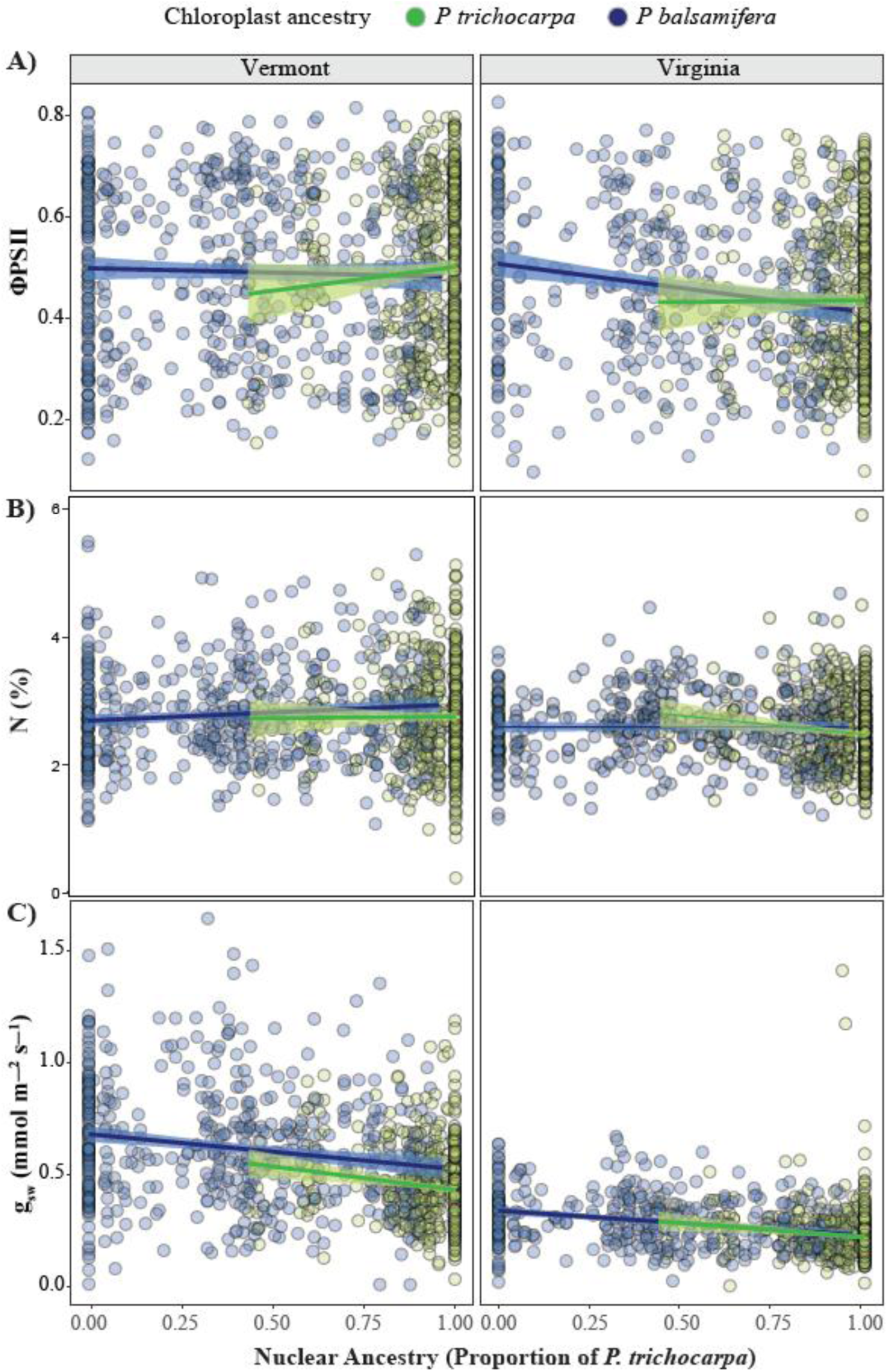
Relationships between A) the quantum efficiency of light in PSII (ΦPSII), B) nitrogen content (N, %), and C) stomata conductance (g_sw_, mmol m^−2^ s^−1^) and the interaction between nuclear genomewide average ancestry and chloroplast ancestry across two environments. Individual data points represent a *Populus* clone assessed within each environment. Regression lines depict the relationship between trait values given estimates of genome wide average ancestry and chloroplasty ancestry with 95% confidence intervals shaded. Colors denote chloroplast ancestry, with green indicating *P. trichocarpa* chlorotypes, and blue representing *P. balsamifera* chlorotypes.

A significant positive effect of cytonuclear interactions on nitrogen content in leaves was detected, with the interaction between nuclear and chloroplast ancestry (*slope* = 0.43, *p* < 0.05) indicating that concordant ancestry can increase nitrogen content (Table 1, Fig. 4B). If genotypes had 50% *P. trichocarpa* nuclear ancestry and the *P. trichocarpa* chlorotype, they exhibited increased nitrogen content relative to those that had the *P. balsamifera* chlorotype. Overall, nitrogen content in leaves was significantly reduced in the Virginia garden relative to the Vermont garden, with an average decrease of 0.24% (*p* < 0.001), indicating site-specific effects. These results suggest that nitrogen content can be impacted by mismatch in cytonuclear ancestry and local site-specific conditions.

While cytonuclear interactions did not influence stomatal conductance (g_sw_), a significant negative effect of both nuclear ancestry and common garden environment was observed (Fig. 4C). g_sw_ decreased at a rate of 0.16 mmol m^-2^ s^-1^ as the nuclear ancestry shifted from *P. balsamifera* towards *P. trichocarpa* (*p* < 0.001, Table 1) reflecting species-specific differences in water-use strategies. Furthermore, g_sw_ was significantly reduced in Virginia relative to Vermont, with an average decrease of 0.26 mmol m⁻² s⁻¹ in the Virginia garden (*p* < 0.001), indicating strong site-specific environmental effects. Chloroplast ancestry did not appear to have a significant effect on g_sw_ (Figure 4C). These results indicate that g_sw_ is influenced by nuclear ancestry and environmental conditions rather than cytonuclear interactions.

## Discussion

The theory of co-adaptation suggests the nuclear and cytoplasmic genomes have co-evolved to optimize physiological processes critical to plant fitness across environments [2,9,10,23,55]. However, hybridization can disrupt co-adapted gene complexes with a range of fitness consequences [22]. Across six contact zones between two sister *Populus* species, we found that nuclear–chloroplast (N-cp) genes often introgress independently of chloroplast ancestry. However, the extent of co-introgression varied by contact zone, suggesting that local environmental conditions may play a stronger role influencing patterns of co-introgression. Further, common garden experiments revealed that chloroplast and nuclear mismatch can modify photosynthetic efficiency, although the effect of the chloroplast-nuclear mismatch can be environmentally dependent. Overall, our results suggest that chloroplast and nuclear genomes interact to influence adaptive trait variation. Given their importance for plant function, understanding how these interactions respond to climate change will be critical for predicting plant performance under future climates.

### Chloroplast divergence and directional introgression are shaped by selection and demography

A comparison of chlorotypes for *P. trichocarpa* and *P. balsamifera* indicated species-specific differences associated with fixed amino acid changes. Despite only ∼2% sequence divergence, nine fixed amino acid changes were identified in genes associated with water-use efficiency (*rpo*C2, *ycf*1, Ruiz-Nieto et al., 2015), high-light oxidative stress (*ndh*D, *ndh*K, Peng et al., 2011; Daniell et al., 2016), cold tolerance (*rps*8, Zhang et al., 2020), splicing and biogenesis (*mat*K, *rpo*A). The maintenance of these species-specific differences suggests that they may underlie functional variation important to adaptive differentiation between the two species.

The geographic distribution of chlorotypes largely followed species-specific geographic distributions. However, we detected a chloroplast capture for *P. balsamifera* in the Alaska contact zone and evidence of *P. balsamifera* expansion into the range of *P. trichocarpa* in the Chilcotin contact zone. While selection may favor chlorotypes due to adaptation or cytonuclear compatibilities, neutral processes, including direction of gene flow or demographic history, can also impact the distribution of chloroplast genomes [19,24,26,59]. The complete absence of pure *P. trichocarpa* chlorotypes in the Alaska contact zone suggests that *P. balsamifera* populations likely established first in this region. This is reflected in the persistence of *P. balsamifera* through the Last Glacial Maximum within northern Alaska and adjacent unglaciated regions of Beringia [36,37]. Following glacial retreat, northward expansion of *P. trichocarpa* populations along the Pacific Northwest coast likely contributed pollen needed to facilitate hybrid formation in Alaska. However, no pure *P. trichocarpa* individuals were observed in this transect. In Chilcotin, demographic processes including seed-mediated gene flow, facilitated by landscape features such as river corridors [34], may be driving the expansion of the *P. balsamifera* chlorotype into historically *P. trichocarpa*-occupied areas. The strong association of the *P. balsamifera* chlorotype with colder and drier climatic conditions relative to *P. trichocarpa* suggests that this chlorotype may provide an adaptive advantage in colder environments. Temperature and moisture availability act as selective forces maintaining certain chloroplast genomes in specific regions [60]. Thus, the contemporary distribution of chloroplast genomes likely results from interactions between historical demography, ongoing gene flow, and environmental selection.

### Co-introgression of chloroplast and nuclear-chloroplast genes results from intrinsic and extrinsic selection

Despite predictions that co-adapted genes should co-introgress following hybridization, [30,61], our results show limited evidence of co-introgression. Co-introgression with the chloroplast genome was observed within the Cassiar contact zone for a subset of N-cp genes. The steep chloroplast cline in Cassiar suggests cytonuclear combinations are likely constrained by selection within this region, potentially due to both intrinsic incompatibilities and extrinsic selection. Co-introgressing genes in Cassiar are primarily associated with RNA processing, proteolysis, photosystem I, ribosomal proteins, and transfer RNAs, suggesting that cytonuclear coordination may be required for these essential chloroplast functions. Consistent with findings in other systems, co-introgression between cytoplasmic and nuclear genomes appears to be restricted to a limited number of genes, typically those critical for organelle performance, rather than occurring broadly across the genome [29,61].

In the Cassiar contact zone, the restricted pattern of co-introgression may reflect climate-driven selection, aligning with steep climate turnover from maritime to boreal conditions across this contact zone [62–65]. Chlorotype distribution in this region was strongly associated with climatic variables, including mean annual temperature and relative humidity, suggesting that environmental gradients may impact introgression. Such patterns are expected when nuclear and organellar genomes are adapted to local environments [30] as environmental gradients can reinforce barriers to gene flow by favoring cytonuclear combinations that optimize organelle functions [16]. Therefore, the limited co-introgression of N-cp genes observed in Cassiar may be shaped by environmental selection acting to maintain cytonuclear compatibility needed for adaptation to local environmental conditions. This contrasts with other contact zones where we did not observe a strong role of climate in shaping the distribution of the chlorotype.

Demographic processes may also play a role influencing cytonuclear introgression, particularly in regions where environmental selection is weak [66]. In the Chilcotin, Jasper, Crowsnest, and Wyoming contact zones, we observed no evidence of co-introgression between chloroplast and N-cp genes and only weak associations between chloroplast ancestry and climate. In these regions, recombination has likely played a major role in breaking down co-inherited ancestry blocks over time, particularly in older contact zones with a repeated history of hybridization [66,67]. This process can decouple chloroplast genomes from nuclear genes that interact with them, weakening any signal of co-introgression even if such associations were present initially. Moreover, repeated cycles of range expansion and contraction driven by climatic oscillations [34] can lead to founder effects and genetic drift. These demographic processes can alter the frequency of specific chloroplast haplotypes or nuclear ancestry blocks by chance, either preserving or eliminating cytonuclear combinations independent of their fitness effects. In such contexts, mismatched cytonuclear combinations may persist not because they confer an adaptive advantage, but because selection against them is weak or absent. Consequently, cytonuclear mismatches may be neutral and maintained within hybrid populations. The fitness consequences of such mismatches may only become apparent under specific environmental stressors, such as drought, nutrient limitation, or temperature extremes [68,69]. Thus, the lack of co-introgression observed in these zones likely reflects the combined influence of neutral demographic processes particularly recombination and historical range dynamics and environment-dependent selection.

While our study provides valuable insights into cytonuclear introgression, this analysis relies on the identification of recognized N-cp genes annotated in the *Arabidopsis* genome [49]. The Cytonuclear Molecular Interactions Reference database for Arabidopsis (CyMIRA) provides the most comprehensive list of cytonuclear interactions. However, it is possible that not all potential N-cp genes in *Populus* were captured in this study. Future research should focus on identifying *Populus*-specific N-cp genes by leveraging admixture mapping to detect regions of the genome associated with chloroplast ancestry. This could be followed by functional validation of candidate genes through expression analyses or transgenic approaches [70]. Furthermore, our study focuses on chloroplast-nuclear interactions, but mitochondrial genomes also interact with the nuclear genome to regulate key cellular processes, including respiration and stress responses [4,7,70,71]. Given that mitochondrial genes are also uniparentally inherited and subject to cytonuclear co-adaptation, future work should explore whether nuclear-mitochondrial interactions exhibit similar patterns of introgression.

### Discordance between nuclear and chloroplast genomes reduce light absorbance

The nuclear and chloroplast genomes interact to influence traits important to adaptation, including physiological traits critical to plant fitness [1,8,14,15]. The efficiency of photosystem II (ΦPSII) and maximum fluorescence (Fm) are particularly important during early tree establishment as reduced light absorbance limits photosynthetic capacity essential for survival and performance [72]. Here, we provide evidence that cytonuclear interactions negatively affect ΦPSII and Fm, with mismatched chloroplast and nuclear ancestry reducing their efficiency in two different environments. On average, individuals with matched chloroplast and nuclear ancestry had higher light absorbance than those with mismatched ancestry. The higher light absorbance for individuals with matched genome ancestry highlights the importance of positive intergenomic epistasis for plant adaptation. For admixed genotypes, those carrying the *P. balsamifera* chlorotype had a slight advantage when nuclear ancestry was evenly mixed (∼50% *P. trichocarpa* or *P. balsamifera*). However, this advantage diminished as *P. trichocarpa* nuclear ancestry increased, suggesting that cytonuclear mismatches reduce the efficiency of light absorbance, particularly in hybrids with a *P. balsamifera* chlorotype and greater proportion of *P. trichocarpa* nuclear genome ancestry. These findings align with previous work in *Arabidopsis* [2], *Ipomopsis* [19], *Helianthus* [15], and wild barley [73], that showed mismatches between nuclear and chloroplast genome ancestry reduced the efficiency of key physiological traits. As the reduction in ΦPSII and Fm was observed in both common gardens, the effect is likely driven by intrinsic incompatibilities rather than an environment-specific stress responses [30,68]. These incompatibilities may reflect Bateson-Dobzhansky-Muller (BDM) incompatibilities, which predict negative fitness outcomes associated with epistatic interactions between divergent alleles [21,22]. This supports the idea that *P. trichocarpa* and *P. balsamifera*, which diverged over ∼2.12 million years ago (Bolte et al., 2024), have evolved lineage-specific cytonuclear interactions. Disruptions of those interactions via hybridization may lead to the evolution of postzygotic barriers, limiting the direction and extent of introgression in hybrid zones.

In contrast to the consistent effects observed for light absorbance, the impact of cytonuclear interactions on nitrogen content in leaves was environment specific. Nitrogen plays an essential role in the photosynthetic processes, contributing to the synthesis of light-harvesting proteins, bioenergetic molecules, and enzymes like Rubisco that are critical for carbon fixation [74]. Matched nuclear and chloroplast ancestry were associated with increased leaf nitrogen content, suggesting that cytonuclear concordance enhances nitrogen-related physiological function beyond its role in light absorbance. The positive effect of matching nuclear and chloroplast ancestry on nitrogen content was more pronounced in the Virginia garden, where overall nitrogen availability was significantly lower. This environmental-dependent response parallels patterns observed in *Ipomopsis* and *Penstemon* hybrids, where reciprocal hybrid performance varied across environments [68,69]. In both systems, the strength and direction of cytoplasmic effects shifted with habitat, indicating that environmental selection likely modulates cytonuclear interactions. Differences in leaf nitrogen content across genotypes may reflect either variation in soil nitrogen availability or physiological efficiency in uptake and assimilation [19]. Virginia and Vermont common gardens differ in soil-use history; the Vermont site was formerly a maize field and likely has higher residual nitrogen. Reduced nitrogen content in mismatched genotypes in Virginia suggests that cytonuclear incompatibilities constrain nitrogen assimilation under environments with different soil conditions. These results suggest that cytonuclear interactions may shape hybrid zone dynamics through environmental selection against mismatched genotypes, thereby influencing patterns of introgression.

Broad-sense heritability (H²) captures the proportion of phenotypic variance explained by genetic factors including additive, dominance, and epistatic effects. H² was low for ΦPSII and Fm for the common gardens despite clear evidence for cytonuclear effects on these traits. This suggests that most of the phenotypic variation in light absorbance traits is attributable to environmental or residual sources such as microclimatic variation within blocks, measurement error, unmodeled genotype-by-environment interactions, or epigenetic effects, rather than genetic differences among genotypes. The low H² values for light absorbance traits may reflect plasticity to environmental heterogeneity within the gardens. This is further supported by the fact that R² marginal values, representing variance explained by fixed effects (ancestry and environment) were much lower than R² conditional values, which include both fixed and random effects (Table 1). This suggests that a substantial portion of trait variation is captured by block effects, and that unmeasured environmental factors or fine-scale variation may play a dominant role in shaping observed trait variation. In contrast, traits such as leaf nitrogen content, which showed a positive and significant cytonuclear interaction, exhibited moderate heritability, suggesting that the beneficial effect of cytonuclear concordance on nitrogen content is at least partially heritable across environments. Other traits such as stomatal conductance (g_sw_) and water use efficiency (δ¹³C) also showed moderate heritability but did not exhibit significant cytonuclear interactions. This indicates that while these traits are influenced by genetics, variation observed is not mainly driven by cytonuclear interactions.

## Conclusion

We evaluated the role of cytonuclear interactions to plant adaptation across environments. Species-specific differences in the chloroplast genome may contribute to functional variation critical for adaptation. While we found some evidence of co-introgression between the chloroplast and N-cp genes, differential patterns of co-introgression across contact zones are likely shaped by demographic or environmental differences among contact zones. However, the degree of match or mismatch between chloroplast and nuclear genome ancestry can influence plant function across environments, with implications for adaptive evolution where sister species hybridize, particularly under climate change. Because cytonuclear interactions may constrain physiological performance, studies assessing the adaptive capacity of species and their hybrids should account for these interactions when evaluating responses to environmental change.

## Acknowledgments

We thank Baxter Worthing for assisting with sampling physiological traits in the Vermont common garden and Sara Klopf and Tommy Phannareth for help with physiological trait sampling in the Virginia common garden. We are also grateful to Kyle Peer, Clay Sawyers, and Deborah Bird from the Virginia Tech Reynolds Homestead Forestry Research Station for support with plant propagation. Additionally, we thank Nadia Garzione for contributions to sampling and processing physiological traits. Finally, we appreciate Constance Bolte, Alayna Mead, and Kyra LoPiccolo from the Hamilton Lab for valuable questions and discussions during lab meetings, which helped shape this work.

## Author contributions

Michelle Zavala-Paez led data collection, research design, data analysis, and manuscript writing. Brianna Sutara assembled chloroplast genomes. Stephen Keller contributed to data collection, data analysis, and manuscript writing. Jason Holliday and Matthew Fitzpatrick contributed to data collection and manuscript writing. Jill Hamilton contributed to data collection, research design, data analysis and manuscript writing.

## Data accessibility

Genomic data used in this study from a previously published study Bolte, et al 2024 are archived on NCBI (PRJNA996882). All the scripts to replicate the analyses will be deposited in Dryad upon acceptance.

## References

1. Moison M, Roux F, Quadrado M, Duval R, Ekovich M, Lê DH, Verzaux M, Budar F. 2010 Cytoplasmic phylogeny and evidence of cyto-nuclear co-adaptation in *Arabidopsis* thaliana. Plant Journal 63, 728–738. (doi:10.1111/j.1365-313X.2010.04275.x)

2. Roux F et al. 2016 Cytonuclear interactions affect adaptive traits of the annual plant Arabidopsis thaliana in the field. Proc Natl Acad Sci U S A 113, 3687–3692. (doi:10.1073/pnas.1520687113)

3. Boussardon C et al. 2019 Novel cytonuclear combinations modify *Arabidopsis thaliana* seed physiology and vigor. Front Plant Sci 10. (doi:10.3389/fpls.2019.00032)

4. Wang N et al. 2022 Pan-mitogenomics reveals the genetic basis of cytonuclear conflicts in citrus hybridization, domestication, and diversification. Proceedings of the National Academy of Sciences 119. (doi:10.1073/pnas.2206076119)

5. Green BR. 2011 Chloroplast genomes of photosynthetic eukaryotes. Plant Journal 66, 34–44. (doi:10.1111/j.1365-313X.2011.04541.x)

6. Daniell H, Lin CS, Yu M, Chang WJ. 2016 Chloroplast genomes: Diversity, evolution, and applications in genetic engineering. Genome Biol. 17. (doi:10.1186/s13059-016-1004-2)

7. Møller IM, Rasmusson AG, Van Aken O. 2021 Plant mitochondria – past, present and future. The Plant Journal 108, 912–959. (doi:10.1111/tpj.15495)

8. Greiner S, Bock R. 2013 Tuning a ménage à trois: Co-evolution and co-adaptation of nuclear and organellar genomes in plants. BioEssays 35, 354–365. (doi:10.1002/bies.201200137)

9. Sloan DB. 2015 Using plants to elucidate the mechanisms of cytonuclear co-evolution. New Phytologist. 205, 1040–1046. (doi:10.1111/nph.12835)

10. Postel Z, Sloan DB, Gallina S, Godé C, Schmitt E, Mangenot S, Drouard L, Varré JS, Touzet P. 2023 The decoupled evolution of the organellar genomes of *Silene nutans* leads to distinct roles in the speciation process. New Phytologist 239, 766–777. (doi:10.1111/nph.18966)

11. Sloan DB, Triant DA, Wu M, Taylor DR. 2014 Cytonuclear interactions and relaxed selection accelerate sequence evolution in organelle ribosomes. Mol Biol Evol 31, 673–682. (doi:10.1093/molbev/mst259)

12. Greiner S, Wang X, Herrmann RG, Rauwolf U, Mayer K, Haberer G, Meurer J. 2008 The complete nucleotide sequences of the 5 genetically distinct plastid genomes of Oenothera, subsection Oenothera: II. A microevolutionary view using bioinformatics and formal genetic data. Mol Biol Evol 25, 2019–2030. (doi:10.1093/molbev/msn149)

13. Zupok A et al. 2021 A photosynthesis operon in the chloroplast genome drives speciation in evening primroses. Plant Cell 33, 2583–2601. (doi:10.1093/plcell/koab155)

14. Moffett AA. 1695 Genetical studies in Acacias iii. Chlorosis in interspecific hybrids. Heredity 20, 609–620.

15. Sambatti JBM, Ortiz-Barrientos D, Baack EJ, Rieseberg LH. 2008 Ecological selection maintains cytonuclear incompatibilities in hybridizing sunflowers. Ecol Lett 11, 1082–1091. (doi:10.1111/j.1461-0248.2008.01224.x)

16. Morales HE, Pavlova A, Amos N, Major R, Kilian A, Greening C, Sunnucks P. 2018 Concordant divergence of mitogenomes and a mitonuclear gene cluster in bird lineages inhabiting different climates. Nat Ecol Evol 2, 1258–1267. (doi:10.1038/s41559-018-0606-3)

17. Camus MF, Moore J, Reuter M. 2020 Nutritional geometry of mitochondrial genetic effects on male fertility. Biol Lett 16. (doi:10.1098/rsbl.2019.0891)

18. Bettinazzi S, Liang J, Rodriguez E, Bonneau M, Holt R, Whitehead B, Dowling D, Lane N, Camus MF. 2024 Assessing the role of mitonuclear interactions on mitochondrial function and organismal fitness in natural Drosophila populations. Evol Lett (doi:10.1093/evlett/qrae043)

19. Wu CA, Campbell DR. 2007 Leaf physiology reflects environmental differences and cytoplasmic background in Ipomopsis (Polemoniaceae) hybrids. Am J Bot 94, 1804–1812. (doi:10.3732/ajb.94.11.1804)

20. Tiffin P, Olson MS, Moyle LC. 2001 Asymmetrical crossing barriers in angiosperms. Proceedings of the Royal Society B: Biological Sciences 268, 861–867. (doi:10.1098/rspb.2000.1578)

21. Dobzhansky T. 1937 Genetic nature of species differences. Am Nat 71, 404–420.

22. Burton RS. 2022 The role of mitonuclear incompatibilities in allopatric speciation. Cellular and Molecular Life Sciences. 79. (doi:10.1007/s00018-021-04059-3)

23. Sloan DB, Havird JC, Sharbrough J. 2017 The on-again, off-again relationship between mitochondrial genomes and species boundaries. Mol Ecol 26, 2212–2236. (doi:10.1111/mec.13959)

24. Lee-Yaw JA, Grassa CJ, Joly S, Andrew RL, Rieseberg LH. 2019 An evaluation of alternative explanations for widespread cytonuclear discordance in annual sunflowers (Helianthus). New Phytologist 221, 515–526. (doi:10.1111/nph.15386)

25. Zhou J et al. 2021 Chloroplast genomes in Populus (Salicaceae): comparisons from an intensively sampled genus reveal dynamic patterns of evolution. Sci Rep 11. (doi:10.1038/s41598-021-88160-4)

26. Bock DG, Andrew RL, Rieseberg LH. 2014 On the adaptive value of cytoplasmic genomes in plants. Mol Ecol. 23, 4899–4911. (doi:10.1111/mec.12920)

27. Hill GE. 2015 Mitonuclear Ecology. Mol Biol Evol 32, 1917–1927. (doi:10.1093/molbev/msv104)

28. Hill GE. 2019 Reconciling the Mitonuclear Compatibility Species Concept with Rampant Mitochondrial Introgression. Integr Comp Biol 59, 912–924. (doi:10.1093/icb/icz019)

29. Nikelski E, Rubtsov AS, Irwin D. 2023 High heterogeneity in genomic differentiation between phenotypically divergent songbirds: a test of mitonuclear co-introgression. Heredity (Edinb) 130, 1–13. (doi:10.1038/s41437-022-00580-8)

30. Burton RS, Pereira RJ, Barreto FS. 2013 Cytonuclear genomic interactions and hybrid breakdown. Annu Rev Ecol Evol Syst. 44, 281–302. (doi:10.1146/annurev-ecolsys-110512-135758)

31. Huang DI, Hefer CA, Kolosova N, Douglas CJ, Cronk QCB. 2014 Whole plastome sequencing reveals deep plastid divergence and cytonuclear discordance between closely related balsam poplars, Populus balsamifera and P. trichocarpa (Salicaceae). New Phytologist 204, 693–703. (doi:10.1111/nph.12956)

32. McKown AD, Guy RD. 2018 Hybrid vigour – Poplars play it cool. Tree Physiol 38, 785–788. (doi:10.1093/treephys/tpy055)

33. Niemczyk M, Hu Y, Thomas BR. 2019 Selection of poplar genotypes for adapting to climate change. Forests 10. (doi:10.3390/f10111041)

34. Bolte CE, Phannareth T, Zavala-Paez M, Sutara BN, Can MF, Fitzpatrick MC, Holliday JA, Keller SR, Hamilton JA. 2024 Genomic insights into hybrid zone formation: The role of climate, landscape, and demography in the emergence of a novel hybrid lineage. Mol Ecol 33. (doi:10.1111/mec.17430)

35. Geraldes A, Farzaneh N, Grassa CJ, Mckown AD, Guy RD, Mansfield SD, Douglas CJ, Cronk QCB. 2014 Landscape genomics of Populus trichocarpa: The role of hybridization, limited gene flow, and natural selection in shaping patterns of population structure. Evolution (N Y) 68, 3260–3280. (doi:10.1111/evo.12497)

36. Levsen ND, Tiffin P, Olson MS. 2012 Pleistocene speciation in the genus populus (salicaceae). Syst Biol 61, 401–412. (doi:10.1093/sysbio/syr120)

37. Breen AL, Murray DF, Olson MS. 2012 Genetic consequences of glacial survival: The late Quaternary history of balsam poplar (Populus balsamifera L.) in North America. J Biogeogr 39, 918–928. (doi:10.1111/j.1365-2699.2011.02657.x)

38. Li H, Handsaker B, Wysoker A, Fennell T, Ruan J, Homer N, Marth G, Abecasis G, Durbin R. 2009 The Sequence Alignment/Map format and SAMtools. Bioinformatics 25, 2078–2079. (doi:10.1093/bioinformatics/btp352)

39. Li H. 2011 A statistical framework for SNP calling, mutation discovery, association mapping and population genetical parameter estimation from sequencing data. Bioinformatics 27, 2987–2993. (doi:10.1093/bioinformatics/btr509)

40. Dierckxsens N, Mardulyn P, Smits G. 2017 NOVOPlasty: De novo assembly of organelle genomes from whole genome data. Nucleic Acids Res 45. (doi:10.1093/nar/gkw955)

41. Kearse M et al. 2012 Geneious Basic: an integrated and extendable desktop software platform for the organization and analysis of sequence data. Bioinformatics 28, 1647–1649.

42. Katoh K, Kuma K, Toh H, Miyata T. 2005 MAFFT version 5: improvement in accuracy of multiple sequence alignment. Nucleic Acids Res 33, 511–518. (doi:10.1093/nar/gki198)

43. Kalyaanamoorthy S, Minh BQ, Wong TKF, von Haeseler A, Jermiin LS. 2017 ModelFinder: fast model selection for accurate phylogenetic estimates. Nat Methods 14, 587.

44. Cingolani P, Platts A, Wang LL, Coon M, Nguyen T, Wang L, Land SJ, Lu X, Ruden DM. 2012 A program for annotating and predicting the effects of single nucleotide polymorphisms, SnpEff. Fly (Austin) 6, 80–92. (doi:10.4161/fly.19695)

45. Wang T, Hamann A, Spittlehouse D, Carroll C. 2016 Locally Downscaled and Spatially Customizable Climate Data for Historical and Future Periods for North America. PLoS One 11, e0156720. (doi:10.1371/journal.pone.0156720)

46. Heinze G, Schemper M. 2002 A solution to the problem of separation in logistic regression. Stat Med 21, 2409–2419. (doi:10.1002/sim.1047)

47. Puhr R, Heinze G, Nold M, Lusa L, Geroldinger A. 2017 Firth’s logistic regression with rare events: accurate effect estimates and predictions? Stat Med 36, 2302–2317. (doi:10.1002/sim.7273)

48. Heinze G, Meinhard Ploner, Daniela Dunkler, Harry Southworth, Maintainer Georg Heinze. 2023 Package ‘logistf’.

49. Forsythe ES, Sharbrough J, Havird JC, Warren JM, Sloan DB, Chaw SM. 2019 CyMIRA: The Cytonuclear Molecular Interactions Reference for Arabidopsis. Genome Biol Evol 11, 2194–2202. (doi:10.1093/gbe/evz144)

50. Dias-Alves T, Mairal J, Blum MGB. 2018 Loter: A software package to infer local ancestry for a wide range of species. Mol Biol Evol 35, 2318–2326. (doi:10.1093/molbev/msy126)

51. Mead A et al. 2025 Variation in responses to temperature across the Populus trichocarpa x P. balsamifera hybrid zone predict increased climate suitability for hybrids. Manuscript in preparation

52. Derryberry EP, Derryberry GE, Maley JM, Brumfield RT. 2014 HZAR: hybrid zone analysis using an R software package. Mol Ecol Resour 14, 652–663. (doi:10.1111/1755-0998.12209)

53. Fetter KC, Nelson DM, Keller SR. 2021 Growth-defense trade-offs masked in unadmixed populations are revealed by hybridization. Evolution (N Y) 75, 1450–1465. (doi:10.1111/evo.14227)

54. Bates D. 2016 lme4: Linear mixed-effects models using Eigen and S4. R package version. 1, 1.

55. Sloan DB, Warren JM, Williams AM, Wu Z, Abdel-Ghany SE, Chicco AJ, Havird JC. 2018 Cytonuclear integration and co-evolution. Nat Rev Genet. 19, 635–648. (doi:10.1038/s41576-018-0035-9)

56. Ruiz-Nieto JE, Aguirre-Mancilla CL, Acosta-Gallegos JA, Raya-Pérez JC, Piedra-Ibarra E, Vázquez-Medrano J, Montero-Tavera V. 2015 Photosynthesis and chloroplast genes are involved in water-use efficiency in common bean. Plant Physiology and Biochemistry 86, 166–173. (doi:10.1016/j.plaphy.2014.11.020)

57. Peng L, Yamamoto H, Shikanai T. 2011 Structure and biogenesis of the chloroplast NAD(P)H dehydrogenase complex. Biochim Biophys Acta Bioenerg. 1807, 945–953. (doi:10.1016/j.bbabio.2010.10.015)

58. Zhang J, Guo Y, Fang Q, Zhu Y, Zhang Y, Liu X, Lin Y, Barkan A, Zhou F. 2020 The PPR-SMR protein ATP4 is required for editing the chloroplast rps8 mRNA in rice and maize. Plant Physiol 184, 2011–2021. (doi:10.1104/pp.20.00849)

59. Postel Z, Touzet P. 2020 Cytonuclear genetic incompatibilities in plant speciation. Plants. 9. (doi:10.3390/plants9040487)

60. Galmés J, Flexas J, Keys AJ, Cifre J, Mitchell RAC, Madgwick PJ, Haslam RP, Medrano H, Parry MAJ. 2005 Rubisco specificity factor tends to be larger in plant species from drier habitats and in species with persistent leaves. Plant Cell Environ 28, 571–579. (doi:10.1111/j.1365-3040.2005.01300.x)

61. Beck EA, Thompson AC, Sharbrough J, Brud E, Llopart A. 2015 Gene flow between Drosophila yakuba and Drosophila santomea in subunit V of cytochrome c oxidase: A potential case of cytonuclear cointrogression. Evolution (N Y) 69, 1973–1986. (doi:10.1111/evo.12718)

62. Hamilton JA, Aitken SN. 2013 Genetic and morphological structure of a spruce hybrid (Picea sitchensis × P. glauca) zone along a climatic gradient. Am J Bot 100, 1651–1662. (doi:10.3732/ajb.1200654)

63. Hamilton JA, Lexer C, Aitken SN. 2013 Genomic and phenotypic architecture of a spruce hybrid zone (Picea sitchensis × P. glauca). Mol Ecol 22, 827–841. (doi:10.1111/mec.12007)

64. Hamilton JA, Lexer C, Aitken SN, Hamilton J. 2013 Differential introgression reveals candidate genes for selection across a spruce (Picea sitchensis X P. glauca) hybrid zone. 927–938.

65. Menon M et al. 2021 Adaptive evolution in a conifer hybrid zone is driven by a mosaic of recently introgressed and background genetic variants. Commun Biol 4, 1–14. (doi:10.1038/s42003-020-01632-7)

66. Menon M et al. 2020 Tracing the footprints of a moving hybrid zone under a demographic history of speciation with gene flow. Evol Appl 13, 195–209. (doi:10.1111/eva.12795)

67. McFarlane SE, Jahner JP, Lindtke D, Buerkle CA, Mandeville EG. 2024 Selection leads to remarkable variability in the outcomes of hybridisation across replicate hybrid zones. Mol Ecol 33. (doi:10.1111/mec.17359)

68. Kimball S, Campbell DR, Lessin C. 2008 Differential performance of reciprocal hybrids in multiple environments. Journal of Ecology 96, 1306–1318. (doi:10.1111/j.1365-2745.2008.01432.x)

69. Campbell DR, Waser NM, Aldridge G, Wu CA. 2008 Lifetime fitness in two generations of Ipomopsis hybrids. Evolution (N Y) 62, 2616–2627. (doi:10.1111/j.1558-5646.2008.00460.x)

70. Lian Q, Li S, Kan S, Liao X, Huang S, Sloan DB, Wu Z. 2024 Association Analysis Provides Insights into Plant Mitonuclear Interactions. Mol Biol Evol 41. (doi:10.1093/molbev/msae028)

71. Leister D. 2005 Genomics-based dissection of the cross-talk of chloroplasts with the nucleus and mitochondria in Arabidopsis. Gene 354, 110–116. (doi:10.1016/j.gene.2005.03.039)

72. Tian Y, Ungerer P, Zhang H, Ruban A V. 2017 Direct impact of the sustained decline in the photosystem II efficiency upon plant productivity at different developmental stages. J Plant Physiol 212, 45–53. (doi:10.1016/j.jplph.2016.10.017)

73. Tiwari LD et al. 2024 Cytonuclear interactions modulate the plasticity of photosynthetic rhythmicity and growth in wild barley. Physiol Plant 176. (doi:10.1111/ppl.14192)

74. Mu X, Chen Y. 2021 The physiological response of photosynthesis to nitrogen deficiency. Plant Physiology and Biochemistry. 158, 76–82. (doi:10.1016/j.plaphy.2020.11.019)

