## Supplementary material for "The role of cytonuclear interactions to plant adaptation across a *Populus* hybrid zone": Suplementary_Material_Cytonuclear_Interactions_Populus.docx

**Supplementary methods**

**Chloroplast Genome Assembly**

Chloroplast genomes were assembled using the seed-extend based assembler NOVOPlasty [1]. Ribulose-bisphosphate carboxylase (rbc*L*) from the *P. trichocarpa* chloroplast genome (NC_009143.1) was used as a seed to initiate genome assembly, allowing accurate retrieval of reads and iterative extension to assemble chloroplast genomes. Assembly parameters for chloroplast genome included identification of cytoplasmic genome type (chloroplast), a genome size (range 140,000 to 170,000 bp,), 33bp minimum overlap for assembly extension, and read length 150 bp. Following assembly, the inverted repeat region positions (IR), small single copy region (SSC) and large single copy region (LSC) were identified and manually checked in Geneious v7.1.4 [2]. Chloroplast genomes were annotated using Geneious v7.1 using the *P. trichocarpa* reference chloroplast genome, exhibiting at least 90% similarity. Following annotation, start and end codon positions for each gene were manually verified. In total, 573 out of 574 individual chloroplast genomes were assembled, with one individual (GPR-14_S50_L001) not assembled due to low sequence coverage (< 10x). In addition, 10 individuals (Table S1-2) were removed due to mis-assembly in the small single copy region (SSC) and low coverage in this region.

### **Geographic clines to compare patterns of introgression between chloroplast and nuclear-chloroplast genes**

Transect-specific geographic distances were estimated using the Haversine formula based on the latitude and longitude for each genotype per contact zone. For each genotype, geographic distance $D_{norm}$ was normalized using the following equation:

$$D_{norm}= \frac{D_{i}- D_{min}}{D_{max}+ D_{min}}$$

Where: $D_{i}$ is geographic distance for genotype *i*, $D_{min}$ is the minimum geographic distance in the contact zone and $D_{max}$ is the maximum geographic distance in the contact zone. This normalization scales geographic distances between 0 and 1, allowing for direct comparisons across contact zones.

Geographic cline parameters, including center and width (1/ maximum slope, Derryberry et al., 2014), were estimated for chloroplast, 282 N-cp genes, and 2,611 non-interacting nuclear genes using the R package hzar [3]. For each group, ten different cline models using different combinations of scaling (free or fixed) and tails (both, left, right, mirrored, or none), in addition to the null model (no cline) were fit for each geographic transect. The combination of scaling and tail parameters were included to ensure geographic clines evaluate the range of ancestry frequency variability across the entire contact zone. For each cline model, a burn-in period of 100,000 iterations was used, followed by 500,000 Monte Carlo iterations to estimate the posterior distributions of the model parameters. Across each contact zone, the best fit cline model was calculated using AICc scores, with the selected model having the lowest AICc value. Cline parameter estimates based on the best fit model were compared across transects for each group. To compare patterns of introgression between N-cp and non-interacting nuclear genes, we estimated the mean cline parameters for non-interacting nuclear genes, representing genome-wide introgression patterns. Bootstrapping with 1000 iterations was used to estimate mean cline parameters (center and width). In each iteration, 234 non-interacting nuclear genes were randomly sampled with replacement, and mean parameters were calculated.

### **Statistical analysis**

To assess whether environmental conditions further modify the interaction between nuclear and chloroplast ancestry in physiological trait variation, each trait was modeled using the following formula:

$y_{ijkl}=\mu+N_{i}+C_{j}+E_{k}+N_{i}*C_{j}+C_{j}*E_{k}+N_{i}*C_{j}*E_{k}+R_{l(k)}+ \varepsilon_{ijkl}$

Where $N_{i}$ is the fixed effect of nuclear ancestry *i*, $C_{j}$ is the fixed effect of chloroplast ancestry *j*, $E_{k}$ is the fixed effect of common garden environment *k* accounting for environmental differences across common garden experiments. $N_{i}*C_{j}$ represent the interaction of nuclear and chloroplast ancestry, $C_{j}*E_{k}$ captures how the effect of chloroplast ancestry varies across common garden experiments and $N_{i}*C_{j}*E_{k}$ test whether the interaction between nuclear and chloroplast ancestry is further modified by environmental conditions. The random effect of block nested within garden $R_{l(k)}$ was included to account for site-specific environmental variation within garden and $\varepsilon_{ijkl}$ is the error term.

**References**

1.	Dierckxsens N, Mardulyn P, Smits G. 2017 NOVOPlasty: De novo assembly of organelle genomes from whole genome data. *Nucleic Acids Res* **45**. (doi:10.1093/nar/gkw955)

2.	Kearse M *et al.* 2012 Geneious Basic: an integrated and extendable desktop software platform for the organization and analysis of sequence data. *Bioinformatics* **28**, 1647–1649.

3.	Derryberry EP, Derryberry GE, Maley JM, Brumfield RT. 2014 HZAR: hybrid zone analysis using an R software package. *Mol Ecol Resour* **14**, 652–663. (doi:10.1111/1755-0998.12209)

**Supplementary figures**

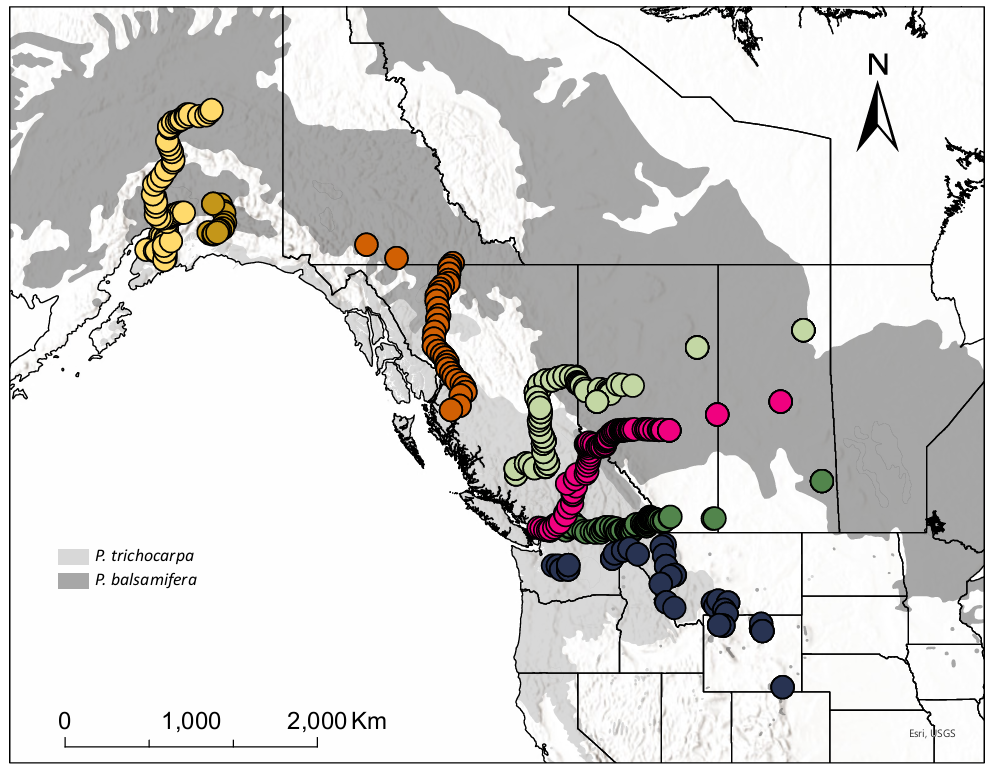

**Figure S1**. Poplar individuals (n = 574) were collected in seven east-west contact zones across the hybrid zone between *P. trichocarpa* and *P. balsamifera*. Contact zones were named Alaska (yellow), Alaska2 (brown), Cassiar (orange), Chilcotin (green), Jasper (pink), Crowsnest (dark green), and Wyoming (dark blue). Light gray represents the distribution range of *P. trichocarpa* and dark gray *P. balsamifera*.

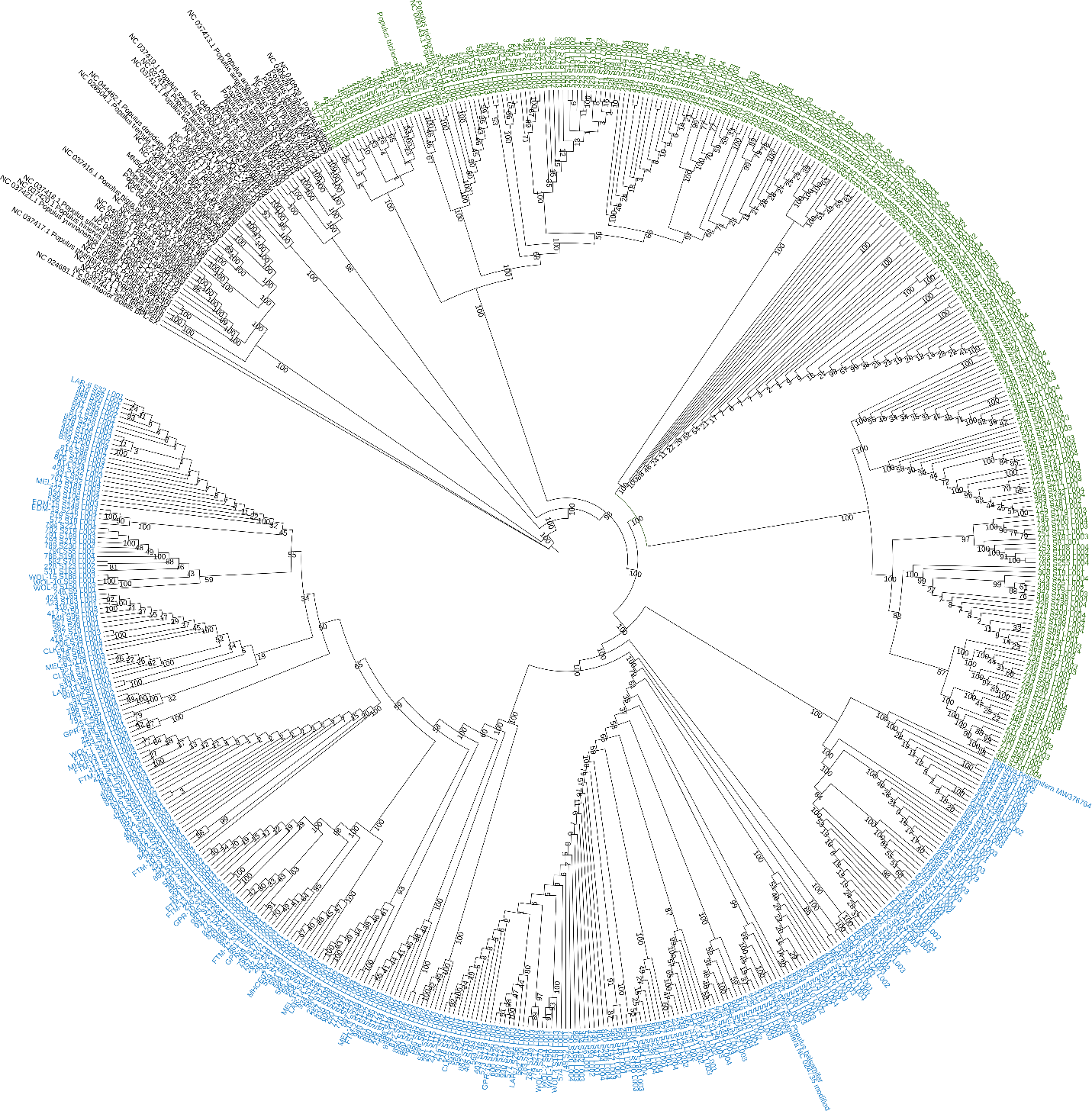

**Figure S2.** Phylogenetic tree of 565 individuals collected from the hybrid zone, along with 58 additional *Populus* and *Salix* species to be used as outgroups. Individuals within the *P. trichocarpa* and *P. balsamifera* clades are highlighted in green and blue, respectively. Branch lengths were not scaled in this representation to facilitate visualization

.
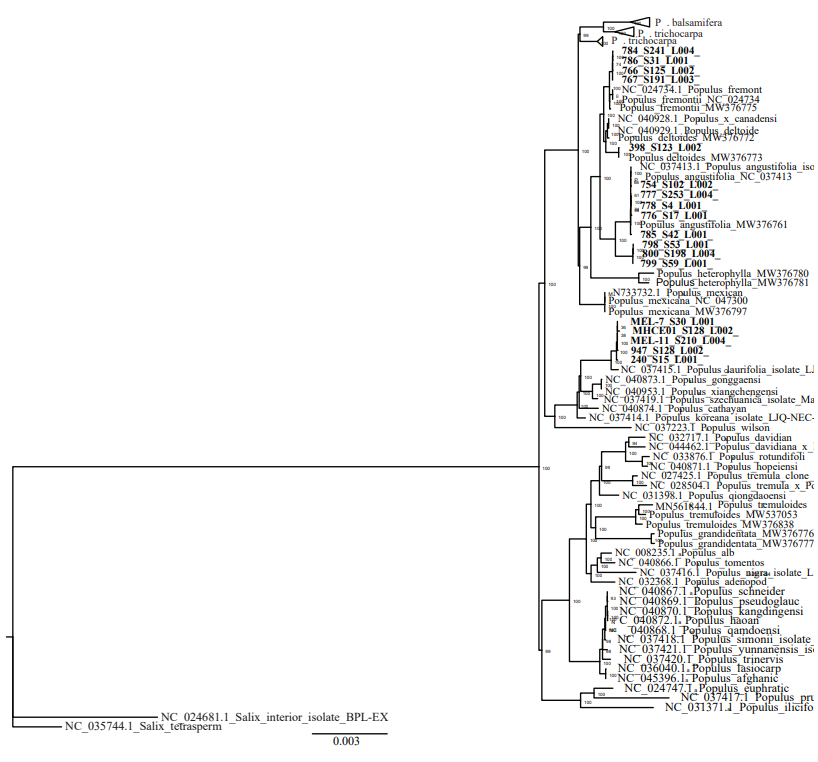

**Figure S3.** Phylogenetic tree of the 565 genotypes collected across the natural hybrid zone between *Populus trichocarpa* (n=247) and *P. balsamifera* (n=300) along with 58 additional genotypes representing diversity across the *Populus* and *Salix* genus. Individuals belonging to *P. trichocarpa* and *P. balsamifera* clades were clustered together and are shown at the top of the tree to facilitate visualization. Individuals belonging to other *Populus* clades (n=18), indicating chloroplast ancestry other than *P. trichocarpa* or *P. balsamifera* are indicated in bold.

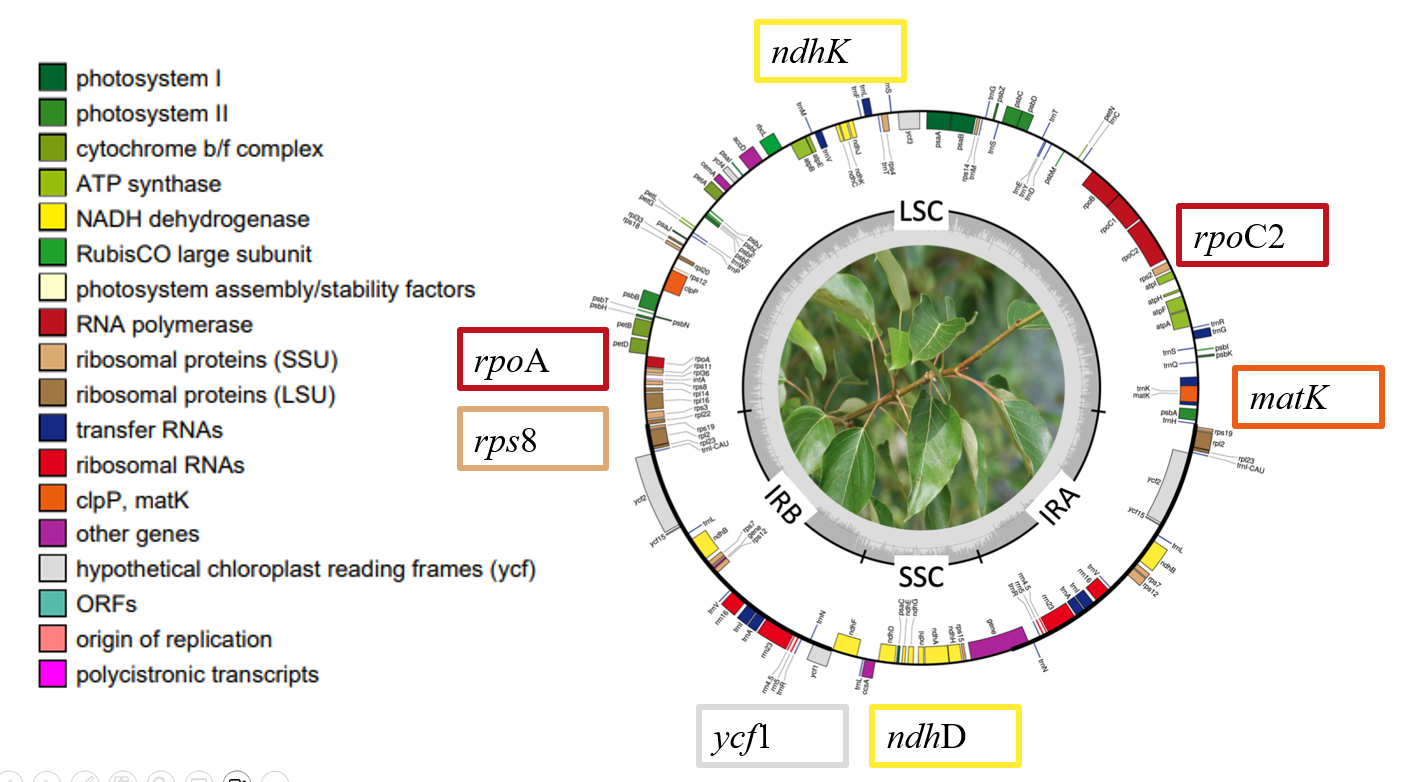

**Figure S4**. *Populus balsamifera* chloroplast genome. Genes with fixed differences between *P. trichocarpa* and *P. balsamifera* chlorotypes are shown in the rectangles. Genes shown on the outside of the circle are transcribed clockwise; genes on the inside are transcribed counterclockwise. The color of the gene boxes indicates the functional group to which the gene belongs. The thick lines indicate the length of the inverted repeats (IRA and IRB), separated by the large single-copy (LSC) and small single-copy (SSC) regions. The dashed darker gray area in the inner circle indicates genome GC content, whereas lighter gray area indicates AT content.

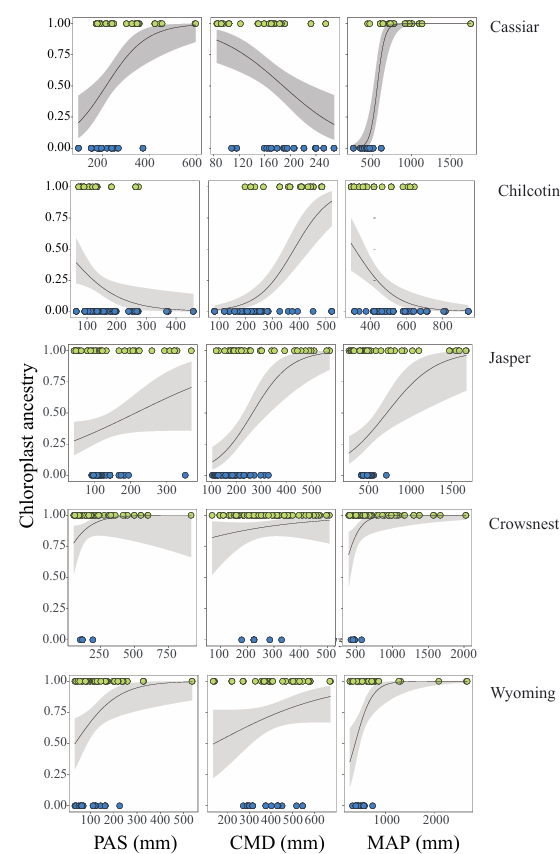

**Figure S5**. A) Relationship between chloroplast ancestry with the precipitation as snow (PAS), climate moisture deficit (CMD), and mean annual precipitation (MAP). Chloroplast ancestry (0 = *P. balsamifera*, 1 = *P. trichocarpa*) is represented as a function of a climate variable. Data points are color-coded by chloroplast ancestry: *P. balsamifera* in blue and *P. trichocarpa* in green. The gray line represents the predicted probabilities of chloroplast ancestry based on the fitted logistic regression, with the shaded blue area indicating 95% confidence intervals for the predictions.

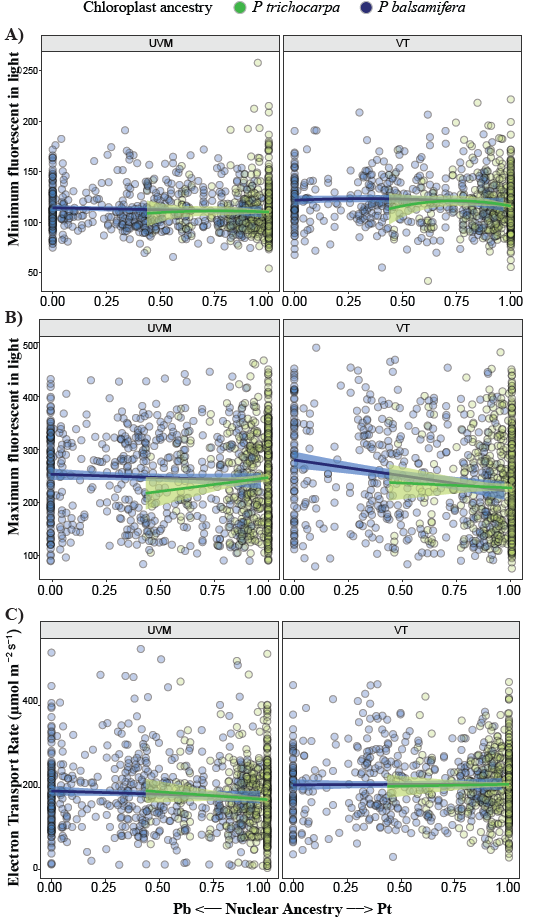

**Figure S6**. A) Relationships between the minimum efficiency in light (Fs), B) maximum efficiency in light (Fm), C) electron transport rate (ETR) and the interaction with nuclear and chloroplast ancestry. Data points represent an individual clone. Lines depict the best fit line for each chlorotype surrounded by 95% confidence shading. Dots and lines are colored coded by chloroplast ancestry, while green indicates *P. trichocarpa* blue represents *P. balsamifera*.
